# Maturation of SARS-CoV-2 Spike-specific memory B cells drives resilience to viral escape

**DOI:** 10.1101/2022.09.30.509852

**Authors:** Roberta Marzi, Jessica Bassi, Chiara Silacci-Fregni, Istvan Bartha, Francesco Muoio, Katja Culap, Nicole Sprugasci, Gloria Lombardo, Christian Saliba, Elisabetta Cameroni, Antonino Cassotta, Jun Siong Low, Alexandra C. Walls, Matthew McCallum, M. Alejandra Tortorici, John E. Bowen, Exequiel A. Dellota, Josh R. Dillen, Nadine Czudnochowski, Laura Pertusini, Tatiana Terrot, Valentino Lepori, Maciej Tarkowski, Agostino Riva, Maira Biggiogero, Alessandra Franzetti Pellanda, Christian Garzoni, Paolo Ferrari, Alessandro Ceschi, Olivier Giannini, Colin Havenar-Daughton, Amalio Telenti, Ann Arvin, Herbert W. Virgin, Federica Sallusto, David Veesler, Antonio Lanzavecchia, Davide Corti, Luca Piccoli

## Abstract

Memory B cells (MBCs) generate rapid antibody responses upon secondary encounter with a pathogen. Here, we investigated the kinetics, avidity and cross-reactivity of serum antibodies and MBCs in 155 SARS-CoV-2 infected and vaccinated individuals over a 16-month timeframe. SARS-CoV-2-specific MBCs and serum antibodies reached steady-state titers with comparable kinetics in infected and vaccinated individuals. Whereas MBCs of infected individuals targeted both pre- and postfusion Spike (S), most vaccine-elicited MBCs were specific for prefusion S, consistent with the use of prefusion-stabilized S in mRNA vaccines. Furthermore, a large fraction of MBCs recognizing postfusion S cross-reacted with human betacoronaviruses. The avidity of MBC-derived and serum antibodies increased over time resulting in enhanced resilience to viral escape by SARS-CoV-2 variants, including Omicron BA.1 and BA.2 sub-lineages, albeit only partially for BA.4 and BA.5 sublineages. Overall, the maturation of high-affinity and broadly-reactive MBCs provides the basis for effective recall responses to future SARS-CoV-2 variants.

## INTRODUCTION

Since its appearance in 2019, severe acute respiratory syndrome coronavirus 2 (SARS-CoV-2) has rapidly spread worldwide resulting in more than 500 million infections and 6.2 million deaths. The virus has evolved into variants of concern (VOC), including the currently circulating Omicron (B.1.1.529) sublineages, which have infected many convalescent and vaccinated individuals (Tegally et al., 2022; Viana et al., 2022). These VOC have accrued several mutations, in particular in the Spike (S), resulting in the reduction or complete loss of neutralizing activity of polyclonal serum and monoclonal antibodies of individuals who were infected or vaccinated with the prototypic SARS-CoV-2 S (Bowen et al., 2022a; Cameroni et al., 2022; Cele et al., 2021; Chen et al., 2021; Collier et al., 2021; Garcia-Beltran et al., 2021; Hoffmann et al., 2020; McCallum et al., 2022; McCallum et al., 2021; Meng et al., 2022; Rees-Spear et al., 2021; Shen et al., 2021; Supasa et al., 2021; Walls et al., 2022; Wang et al., 2021a; Wang et al., 2022; Zhou et al., 2021). Besides serum antibodies, memory B cells (MBCs) induced by infection or vaccination play a major role in humoral immunity through recall responses to a second encounter with the same or a related pathogen. Although several studies reported a progressive decline of serum antibody titers over time in convalescent and vaccinated individuals, SARS-CoV-2-specific MBCs have been shown to increase or remain stable in number and to produce neutralizing antibodies (Dan et al., 2021; Gaebler et al., 2021; Goel et al., 2021a; Goel et al., 2021b; Rodda et al., 2021; Roltgen et al., 2020; Sokal et al., 2021b; Walls *et al*., 2022).

In this study, we performed single-MBC repertoire analysis from longitudinal samples of convalescent or healthy individuals receiving up to three doses of the Pfizer/BioNtech BNT162b2 mRNA vaccine. We found that, while SARS-CoV-2-specific serum antibodies waned, MBCs increased progressively in frequency and avidity, reaching steady-state levels that remained stable up to 16 months after infection or after the third vaccine dose. Vaccination-induced MBCs mainly targeted the prefusion SARS-CoV-2 S conformation, while infection-induced MBCs recognized also the postfusion conformation and cross-reacted with the S of human betacoronaviruses HCoV-HKU1 and HCoV-OC43. We show that the increased avidity of MBC-derived antibodies provides a mechanism of resilience against emerging variants, including Omicron BA.1 and BA.2 sublineages.

## RESULTS

### Progressive maturation of MBCs and serum antibodies following SARS-CoV-2 infection

Blood samples were collected from 64 individuals diagnosed with COVID-19 between March and November 2020 and from 13 individuals diagnosed between December 2020 and January 2021 after an outbreak of the SARS-CoV-2 Alpha variant (**Table S1**). Peripheral blood mononuclear cells (PBMCs) were isolated for antigen-specific memory B cell repertoire analysis (AMBRA) (Pinna et al., 2009) (**Figure 1A**). PBMCs were stimulated in multiple cultures with IL-2 and the Toll-like receptor 7/8-agonist R848 to promote the selective proliferation and differentiation of MBCs into antibody-secreting cells. The culture supernatants were collected on day 10 and screened in parallel using multiple ELISA assays to detect antibodies of different specificities and to determine the frequencies of antigen-specific MBCs expressed as a fraction of total IgG^+^ MBCs (**Figures S1A-S1C**).

**Figure 1.**
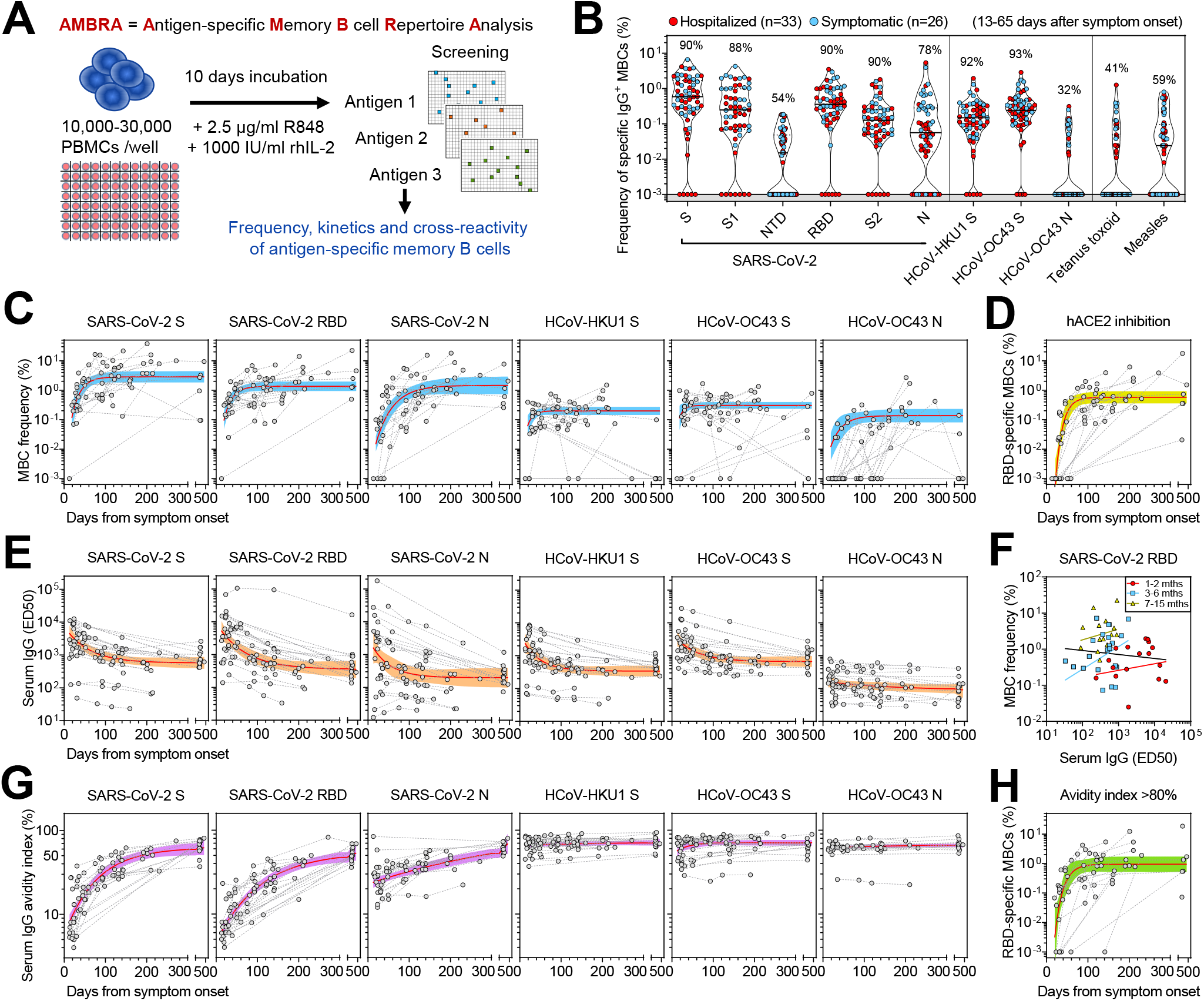
Early response, RBD immunodominance, kinetics and affinity maturation of memory B cells primed by Wuhan SARS-CoV-2. (A) Scheme of the AMBRA method used in this study. PBMC, peripheral blood mononuclear cells. R848, agonist of Toll-like receptors 7 and 8. rhIL2, recombinant human interleukin 2. (B) Frequency of SARS-CoV-2-specific MBCs isolated between 13 and 65 days after symptom onset from n = 59 donors (33 hospitalized, red, and 26 symptomatic, blue) after the analysis of 5,664 MBC cultures. Shown is the reactivity to antigens of SARS-CoV-2 and other betacoronaviruses (HCoV-HKU1 and HCoV-OC43): Spike (S), S1 domain, N-terminal domain (NTD), receptor-binding domain (RBD), S2 domain, Nucleoprotein (N). Reactivities to Tetanus toxoid and to Measles virus (lysate) are included as controls. Median and quartiles are shown as plain and dotted lines, respectively. Percentages of donors with detectable specific MBCs are indicated above each set of data. (C) Frequency of MBCs specific for SARS-CoV-2 S, RBD and N, HCoV-HKU1 S, HCoV-OC43 S and N from n = 23 donors followed-up up to 469 days after symptom onset. Frequencies were obtained from the analysis of 6,336 MBC cultures (66 samples, minimum 2 samples per donor). Black dotted lines connect samples from the same donor. A one-phase association kinetics model (red line) was calculated from all the non-null values of each sample. The area within 95% confidence bands is shown in blue. (D) Frequency of SARS-CoV-2 RBD-specific MBC producing antibodies showing inhibition of RBD binding to ACE2 from n = 23 donors. A one-phase association kinetics model (red line) was calculated from all the non-null values of each sample and the area within 95% confidence bands is shown in yellow. (E) Serum IgG ED50 titers to SARS-CoV-2 S, RBD and N, HCoV-HKU1 S, HCoV-OC43 S and N of samples collected from 29 donors analyzed up to 469 days after symptom onset. A one-phase decay kinetics model (red line) was calculated from all the non-null values of each sample and the area within 95% confidence bands is shown in orange. (F) Correlation analysis between frequency of SARS-CoV-2 RBD-specific MBCs and serum RBD-specific IgG titers of n = 56 samples from n = 18 donors collected at different time points. All samples (black line): Spearman r = -0.102 (95% confidence interval -0.363 to 0.173; non-significant P = 0.45). Samples at 1-2 months (n = 18, red line): Spearman r = 0.112 (95% confidence interval -0.387 to 0.561; non-significant P = 0.66). Samples at 3-6 months (n = 23, blue line): Spearman r = 0.214 (95% confidence interval -0.229 to 0.584; non-significant P = 0.33). Samples at 7-15 months (n = 15, yellow line): Spearman r = 0.221 (95% confidence interval -0.343 to 0.668; non-significant P = 0.43). (G) Serum IgG avidity indexes to SARS-CoV-2 S, RBD and N, HCoV-HKU1 S, HCoV-OC43 S and N of samples collected from 29 donors analyzed up to 469 days after symptom onset. A one-phase association kinetics model (red line) was calculated from all the non-null values of each sample and the area within 95% confidence bands is shown in violet. (H) Frequency of SARS-CoV-2 RBD-specific B cells with an avidity index greater than 80%. A one-phase association kinetics model (red line) was calculated from all the non-null values of each sample and the area within 95% confidence bands is shown in green.

Repertoire analysis of MBCs collected at early time points after infection with SARS-CoV-2 Wuhan-Hu-1 (13-65 days after symptom onset) showed that 90% of the donors had detectable MBCs specific for the prefusion-stabilized SARS-CoV-2 S ectodomain trimer (Walls et al., 2020) (**Figures 1B** and **S1D**). The magnitudes of MBC responses were heterogenous with frequencies ranging between 0 and 6.6% for S-specific MBCs across individuals. In most cases RBD-specific MBCs dominated the response to S, whereas NTD- and S_2_-specific MBCs were present at lower frequencies, concurring with the fact that most mAbs cloned from the memory B cells of previously infected subjects target the RBD (McCallum *et al*., 2021; Piccoli et al., 2020). Overall, S-specific MBCs were present at higher frequencies than N-specific MBCs (median: 0.59% vs 0.06%, respectively). A similar magnitude and reactivity of the MBC response was observed in the individuals infected with the SARS-CoV-2 Alpha variant, with a higher frequency of MBCs specific for RBD carrying the N501Y mutation as compared to the Wuhan-Hu-1 RBD (**Figure S1E**).

By analyzing longitudinal samples collected up to 16 months after infection, we found that S-, RBD-, NTD-, S_2_- and N-specific MBCs progressively rose in frequency in the first 6-8 months, reaching up to 20% of total IgG MBCs in some cases, followed by a plateau (Dan *et al*., 2021). Conversely, the frequency of MBCs specific for the S and N proteins of common cold coronaviruses remained largely constant over time (**Figure 1C** and **S1F**). The expansion of RBD-specific MBCs was accompanied by an increase of MBCs producing antibodies that blocked RBD binding to human ACE2, a correlate of neutralization (Piccoli *et al*., 2020) (**Figures 1D** and **S1G**).

Analysis of longitudinal serum samples showed that IgG antibodies to SARS-CoV-2 S, RBD and N progressively decreased and reached a plateau, which paralleled the rise of MBCs, consistent with previous reports (Achiron et al., 2021; Gaebler *et al*., 2021; Khoury et al., 2021) (**Figure 1E**). We observed a modest increase of IgG antibodies recognizing HCoV-HKU1 and HCoV-OC43 S early after SARS-CoV-2 infection, which rapidly dropped to levels maintained until the end of the observation period (**Figure 1E**). No correlation was observed between serum IgG S antibody levels and MBC frequencies in the subjects tested with both assays over the 500-day period analyzed (**Figure 1F**).

The binding avidity of serum- and MBC-derived specific antibodies was expressed as an avidity index by measuring antibody binding in presence of a chaotropic agent (**Figure S1H**). The avidity of serum IgG antibodies to SARS-CoV-2 S, RBD and N increased over time reaching levels comparable to those observed for serum IgG antibodies to HKU1 and OC43 antigens (**Figure 1G**). Similarly, the frequency of high-avidity MBC-derived RBD-specific antibodies increased over time reaching a plateau after approximately 3-to-4 months after infection (**Figure 1H**).

The rapid decline of serum antibodies is consistent with an initial generation of many short-lived plasma cells from either naïve B cells or cross-reactive MBCs. However, the clonal analysis of serial samples reveals a rapid expansion of S- and RBD-specific MBCs followed by a progressive maturation consistent with a germinal center reaction leading to the generation of plasma cells and MBCs with increased affinity.

### Three mRNA-vaccine doses induce high-avidity MBCs and serum antibodies

The frequency and fine specificity of MBCs were analyzed after the first, second and third administration of the Pfizer/BioNtech BNT162b2 mRNA vaccine in two cohorts of healthy individuals. Vaccine recipients were either naïve (n=46) or immune (n=32) to SARS-CoV-2 due to infection occurring 53-389 days before the first dose. In most naïve individuals, the first vaccine dose induced SARS-CoV-2 S-specific MBCs at frequencies comparable to those found in samples collected from convalescent individuals at similar timepoints post antigen exposure (**Figure 2A**). The second and third doses resulted in a further 5-fold and 10-fold increase of median MBC frequency, respectively (**Figure 2A**). As expected, vaccination of naïve donors did not elicit N-specific MBCs, the few exceptions likely reflecting cross-reactivity with other betacoronaviruses (**Figure S2A**). Remarkably, the first vaccine dose induced very high MBC S-specific frequencies in previously infected donors, exceeding by ∼40-fold those found in naïve vaccinated or convalescent individuals. Additional vaccine doses did not result in further MBC increase and did not alter the frequency of MBCs specific for the S of the common cold coronaviruses HCoV-HKU1 and HCoV-OC43 (**Figures 2B** and **S2B**).

**Figure 2.**
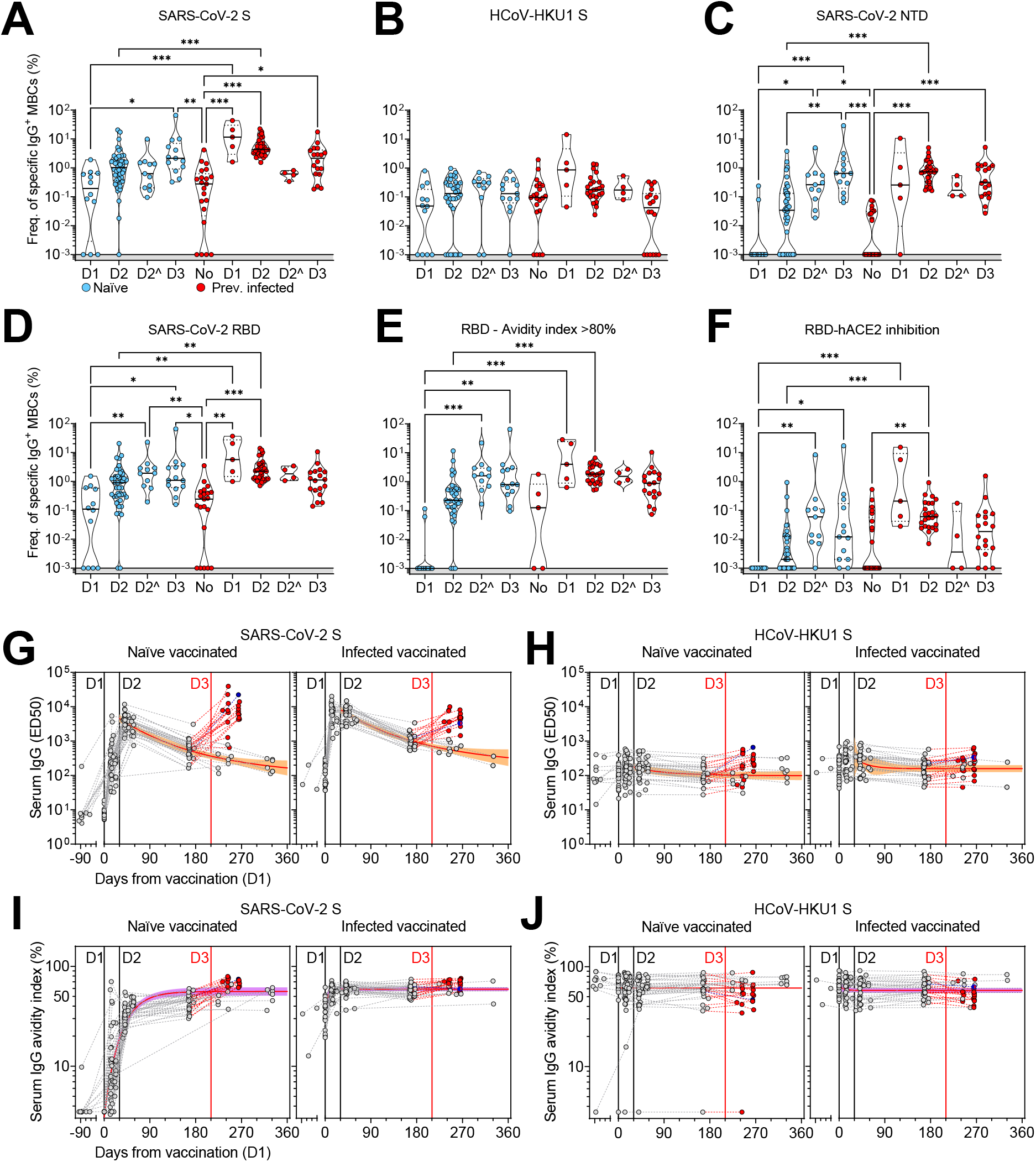
Characterization of vaccine-induced MBC- and serum-derived antibody response in naïve and SARS-CoV-2 immune donors. (A-D) Frequency of MBCs specific for SARS-CoV-2 S (A), HCoV-HKU1 S (B), SARS-CoV-2 NTD (C) and RBD (D) of n = 12, 45 and 13 naïve donors and 5, 31, and 18 previously infected donors 10-35 days after the first (D1), the second (D2) and the third dose (D3) of Pfizer/BioNtech BNT162b2 mRNA vaccine, respectively. Shown are also 11 naïve and 4 immune donors whose MBCs were isolated 125-293 days after the second dose (D2^). Median frequencies are compared withing donor groups and between respective vaccine doses as well as to a group of n = 21 convalescent donors at 18-30 days after symptom onset. Significant differences are indicated as *** (p-value < 0.001); ** (p < 0.002), * (p < 0.033), ns (non-significant, p > 0.12). (E) Frequency of SARS-CoV-2 RBD-specific MBCs with an avidity index greater than 80%. (F) Frequency of SARS-CoV-2 RBD-specific MBCs inhibiting binding of RBD to ACE2. (G-H) Serum IgG ED50 titers to SARS-CoV-2 S (G) and HCoV-HKU1 S (H) of samples collected from n = 47 naïve (left) and 32 immune donors (right) 10-35 days after the first (D1), the second (D2) and the third dose (D3) of Pfizer/BioNtech BNT162b2 mRNA vaccine, respectively. A one-phase decay kinetics model (red line) was calculated from all the non-null values of each sample and the area within 95% confidence bands is shown in orange. 37 samples collected from individuals who received a third dose (red) or had a second SARS-CoV-2 infection (blue) were excluded from the decay analysis. (I-J) Serum IgG avidity indexes to SARS-CoV-2 S (I) and HCoV-HKU1 S (J) of the same samples shown in panels G-H. A one-phase association kinetics model (red line) was calculated from all the non-null values of each sample and the area within 95% confidence bands is shown in violet.

After administration of two vaccine doses, the response in both naïve and infected donors was dominated by RBD-specific MBCs, while MBCs specific for the NTD or the S_2_ subunit were present at low to undetectable levels (**Figures 2C, 2D** and **S2C**). Interestingly, NTD-specific, but not S2-, MBCs increased over time in naïve donors (**Figure 2C** and **S2C**). High-avidity RBD-specific and ACE2-blocking antibodies were detected only after the second dose in naïve individuals, whereas these antibodies were detected after one dose in all the previously infected donors and were not further boosted upon subsequent immunizations (**Figure 2E** and **2F**).

When we analyzed longitudinal samples of serum antibodies against SARS-CoV-2 S and RBD, we observed that a single immunization induced highly heterogenous antibody levels in naïve donors that were further increased in all samples after the second and the third immunization (**Figures 2G** and **S2D**). In infected donors, the titers of serum antibodies had reached the maximal level after the first immunization, with no further increase after the second or the third immunization. The serum titers of S-specific antibodies declined similarly over time in naïve and infected donors with a half-life of 4 months. As expected, no overall variation in N-specific antibody titers was observed (**Figure S2E**). While the avidity of SARS-CoV-2 S- and RBD-specific serum antibodies in naïve donors rapidly increased after vaccination, the avidity in infected donors was found to be high before vaccination with no further increase over time (**Figures 2I, S2F** and **S2G**). In both naïve and infected donors, vaccination did not increase the presence or avidity of antibodies specific for the S of the human betacoronaviruses HCoV-HKU1 and HCoV-OC43, concurring with recent data (Walls *et al*., 2022) (**Figures 2H, 2J, S2H** and **S2I**).

Collectively, these findings indicate that, while in infected donors a single dose of an mRNA vaccine is sufficient to boost a high-avidity antibody response to SARS-CoV-2 S, in naïve donors such response is elicited only upon three rounds of immunization.

### The antibody response in naïve vaccinated individuals is skewed towards prefusion SARS-CoV-2 S

The Pfizer/BioNtech BNT162b2 mRNA vaccine was designed to express the full-length SARS-CoV-2 S stabilized in its prefusion conformation through the 2P mutations (Vogel et al., 2021; Wrapp et al., 2020) and recent data suggest that vaccination induce high titers of prefusion S-specific plasma antibodies (Bowen et al., 2021). To assess whether vaccination also induced a MBC response preferentially targeting the prefusion conformation of S as compared to that elicited by natural infection, we analyzed the MBC-derived antibodies for their binding to either the prefusion-stabilized SARS-CoV-2 S, postfusion S2, or both in cohorts of convalescent donors and in naïve or infected vaccinated individuals (**Figure 3A**). We found a strong correlation between antibodies binding to postfusion S_2_ and a structurally-validated postfusion SARS-CoV-2 S_2_ (Bowen *et al*., 2021), thus supporting the use of S_2_ as a proxy for the postfusion conformation of S (**Figure S3)**. Across all individuals, most MBCs induced by natural infection and/or vaccination were specific for the prefusion conformation (**Figure 3B-D**). However, while convalescent and infected vaccinated individuals had 10% and 5% of their MBCs specific for S_2_, respectively, only 1% of MBCs from naïve vaccinated donors were postfusion S_2_-specific. A fraction of MBCs recognized S epitopes shared between prefusion and postfusion conformations. Collectively, these data show that mRNA vaccines primarily induce MBC responses skewed to the prefusion conformation of SARS-CoV-2 S.

**Figure 3.**
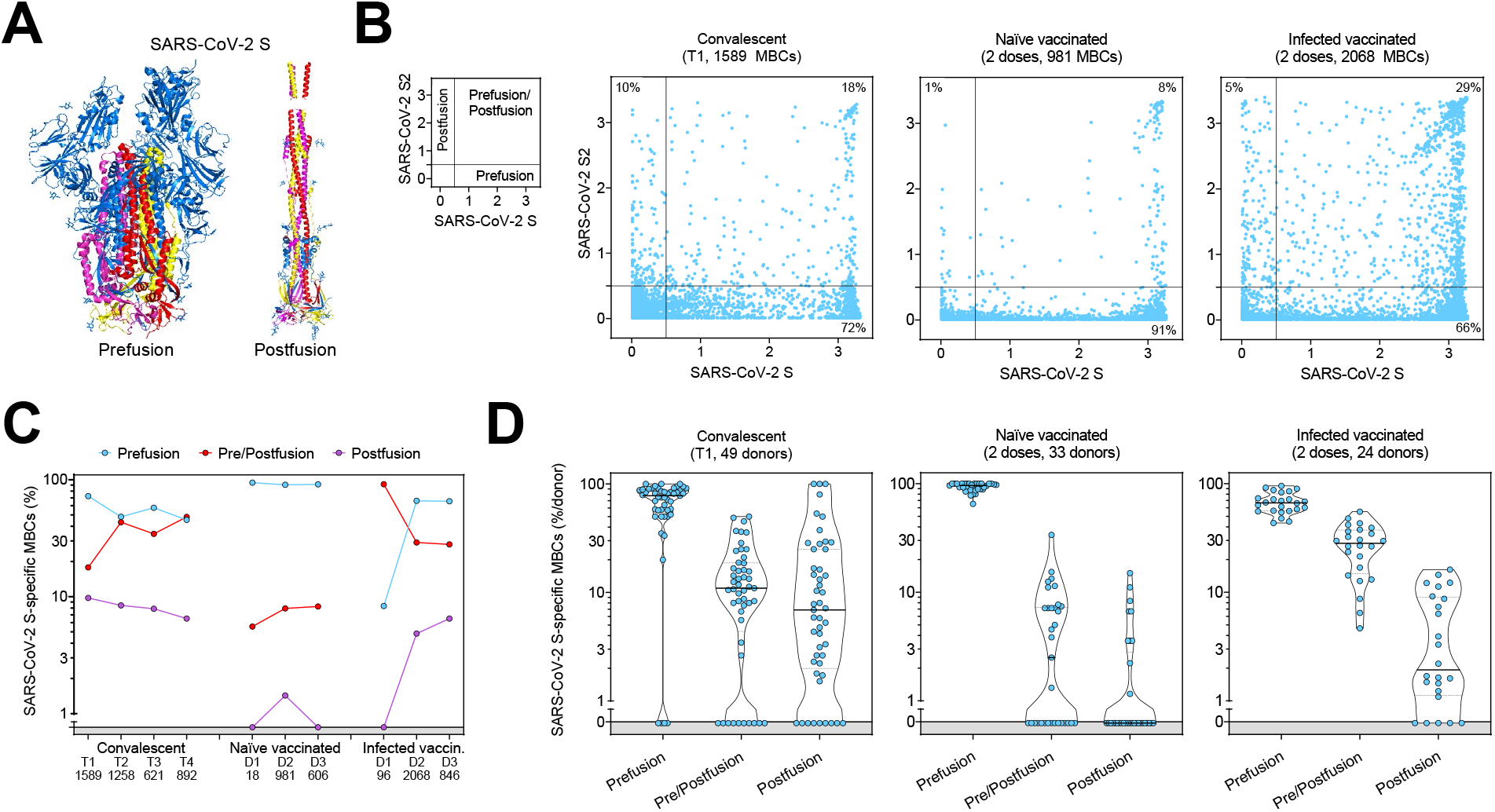
Comparison of the prefusion and postfusion S-specific MBC responses after vaccination or natural infection. (A) Structural representation of SARS-CoV-2 S in its prefusion and postfusion conformation (adapted from PBD 7tat and 7e9t). The three S2 domains that are maintained in both conformations are colored in red, yellow and pink. (B) MBC cross-reactivity between SARS-CoV-2 S (prefusion S) and S2 (postfusion S). Shown are average OD values as measured by ELISA with blank subtracted from n = 2 replicates of 1589, 981 and 2068 MBC cultures analyzed from 49 convalescent, 33 naïve and 24 infected vaccinated donors. Cumulative fraction of MBCs reactive to either prefusion or postfusion S or both is indicated as percentage in the respective quadrant. The small panel on the left describes the distribution of prefusion and/or postfusion S-specific MBCs in the different quadrants. (C) Cumulative fraction of S-specific MBCs reactive to prefusion and/or postfusion S at different timepoints after natural infection (T1, T2, T3 and T4) or vaccine doses (D1, D2, D3). (D) Individual fractions of S-specific MBCs reactive to prefusion and/or postfusion S in 49 convalescent, 33 naïve and 24 infected vaccinated donors.

### Infection- and vaccine-induced MBCs show variable degrees of cross-reactivity against other betacoronaviruses

Unlike serological analyses, the AMBRA method is suitable to dissect the cross-reactivity of individual MBC-derived antibodies, at the single clone level, against a panel of human and animal betacoronaviruses (Forster et al., 2020). Analysis of MBCs from SARS-CoV-2-convalescent and from naïve or infected donors receiving two vaccine doses revealed a high frequency (35-52%) of SARS-CoV-2 S-specific MBCs that cross-reacted with SARS-CoV S, consistent with the high level of S sequence similarity with SARS-CoV-2 (**Figures 4A** and **4B**). Conversely, we observed a low frequency (2-19%) of SARS-CoV-2 S-specific MBCs that cross-reacted with the more divergent S of MERS-CoV, HCoV-HKU1 and HCoV-OC43 betacoronaviruses. A similar trend was observed at late time points in convalescent donors as well as after the third vaccine dose in vaccinated individuals (**Figure S4**). Deconvolution of SARS-CoV-2 S-reactivity revealed a higher frequency of cross-reactive antibodies among the subset of S_2_-specific MBCs (**Figures 4A** and **4B**).

**Figure 4.**
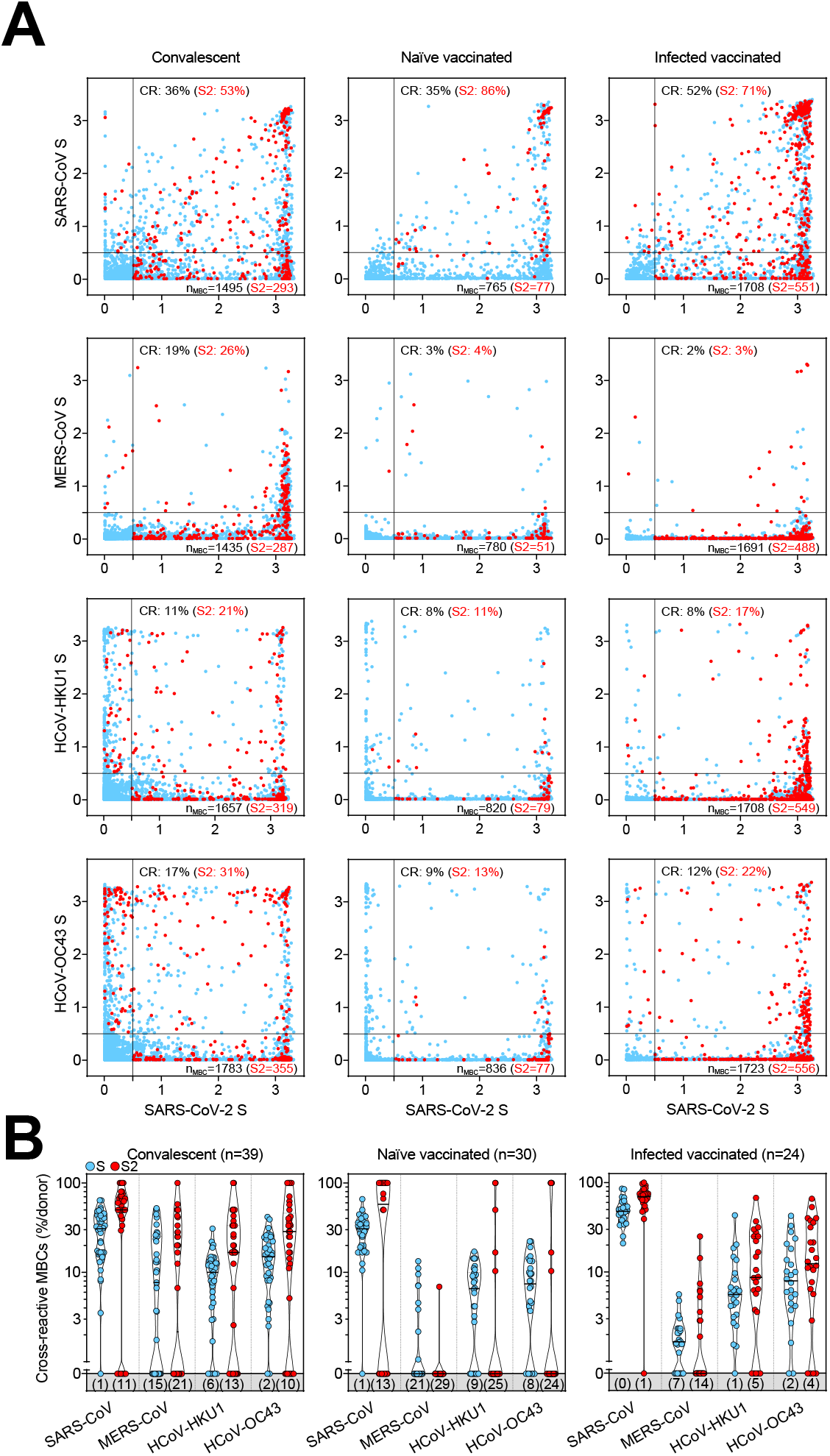
Cross-reactivity to betacoronaviruses of MBCs primed by SARS-CoV-2 infection and/or vaccination. (A) Cumulative MBC cross-reactivity between SARS-CoV-2 S and the four betacoronaviruses SARS-CoV, MERS-CoV, HCoV-HKU1 and HCoV-OC43. Shown are average OD values as measured by ELISA with blank subtracted from n = 2 replicates of 3744, 2880 and 2304 MBC cultures analyzed from 39 convalescent, 30 naïve and 24 infected donors after two vaccine doses. S2-specific MBCs are shown in red. Numbers of S- and S2-specific MBCs are indicated in the bottom-right quadrants of each panel. Cumulative fractions of S- and S2-cross-reactive (CR) MBCs are indicated as percentage in the top-right quadrant. (B) Individual fractions of SARS-CoV-2 S-specific MBCs that cross-react with the four betacoronaviruses in convalescent and vaccinated donors. Numbers in brackets indicate the donors with MBCs showing no cross-reactivity for the respective betacoronavirus S.

The same analysis was performed on RBDs of sarbecoviruses representative of clades 1a (SARS-CoV), 1b (Pangolin Guangxi), 2 (bat ZC45) and 3 (bat BM48-31/BGR/2008) comparing the reactivity of MBCs from SARS-CoV-2-convalescent and from naïve or infected donors receiving two vaccine doses. We found that the frequencies of antibodies cross-reactive to heterologous sarbecovirus RBDs, including those inhibiting binding of RBD to human ACE2, were progressively lower as a function of the decreasing sequence identity with SARS-CoV-2 (**Figure 5A, 5B, S5A** and **S5B**). Infected vaccinees had the highest frequency of cross-reactive MBCs against diverse RBDs, suggesting that increased avidity resulting from multiple and diverse antigenic stimulations may have also contributed to broadening the reactivity towards heterologous sarbecoviruses.

**Figure 5.**
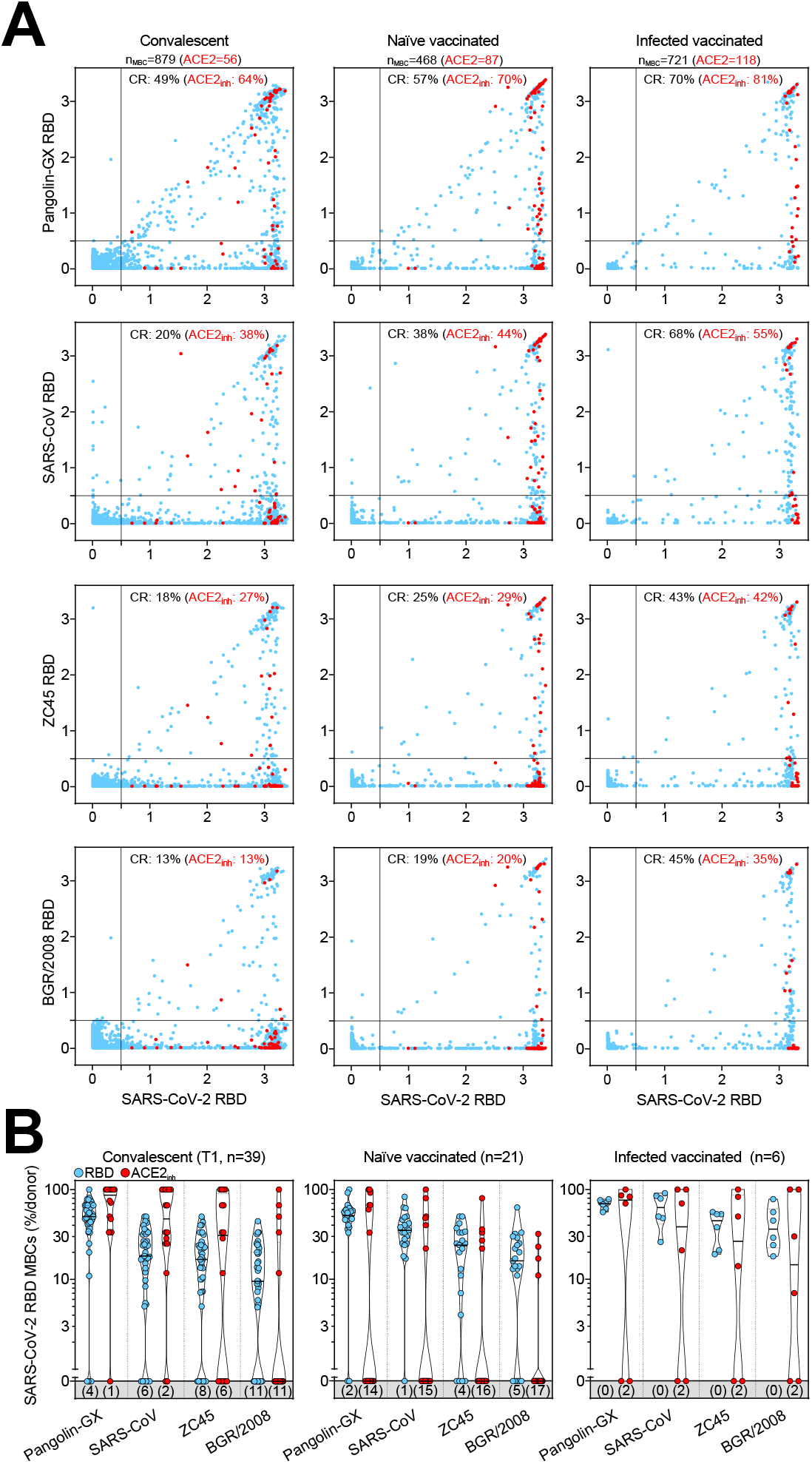
Cross-reactivity to sarbecoviruses of MBCs primed by SARS-CoV-2 infection and/or vaccination. (A) Cumulative MBC cross-reactivity between SARS-CoV-2 RBD and four sarbecoviruses representative of clades 1a (SARS-CoV), 1b (Pangolin Guangxi), 2 (ZC45) and 3 (BM48-31/BGR/2008). Shown are average OD values as measured by ELISA with blank subtracted from n = 2 replicates of 3744, 2016 and 576 MBC cultures analyzed from 39 convalescent donors at two different timepoints and from 21 naïve and 6 infected donors after receiving two vaccine doses. RBD-specific MBCs showing inhibition of binding to ACE2 are shown in red. Cumulative fractions of total and ACE2-inhibiting RBD-cross-reactive (CR) MBCs are indicated as percentage in the top-right quadrant. (B) Individual fractions of SARS-CoV-2 RBD-specific MBCs that cross-react with the four representative sarbecoviruses in convalescent and vaccinated donors. Numbers in brackets indicate the donors with MBCs showing no cross-reactivity for the respective sarbecovirus RBD.

### Affinity maturation of RBD-specific MBCs leads to resilience to viral escape by SARS-CoV-2 variants of concern

In view of the continuing emergence of SARS-CoV-2 VOC, we analyzed at a clonal level the extent to which MBCs elicited by infection with Wuhan-Hu-1-related viruses in early 2020 or by mRNA vaccines could cross-react with RBDs of progressively diverging VOC. We therefore tested MBC-derived antibodies for their ability to retain binding (here measured as less than 2-fold loss compared to Wuhan-Hu-1) to RBD of different VOC, including Beta (B.1.351), Delta (B.1.617.2) and Omicron (B.1.1.529) BA.1, BA.2, BA.3 and BA.4/BA.5 sub-lineages (**Figures 6A, 6B** and **S6A**). In naïve or infected individuals receiving two doses of the Pfizer/BioNtech BNT162b2 mRNA vaccine, mutations in the VOC RBDs were poorly tolerated with the greatest loss of binding observed against Omicron sublineages. Importantly, the third vaccine dose substantially increased the resilience to VOC escape from binding antibodies in both naïve and infected individuals to a degree similar to that observed in convalescent individuals more than one year after infection, in line with analysis of neutralizing antibody responses (Bowen et al., 2022b; Cameroni *et al*., 2022; Park et al., 2022; Walls *et al*., 2022). Interestingly, in individuals given three vaccine doses, the higher fraction of MBCs cross-reactive with VOC RBDs was characterized by high-avidity and ACE2-blocking activity (**Figures 6A, 6B, S6A** and **S6B**). However, in all cohorts analyzed, the binding of MBC-derived antibodies was substantially reduced when tested against BA.4/BA.5 RBDs (**Figures 6A**).

**Figure 6.**
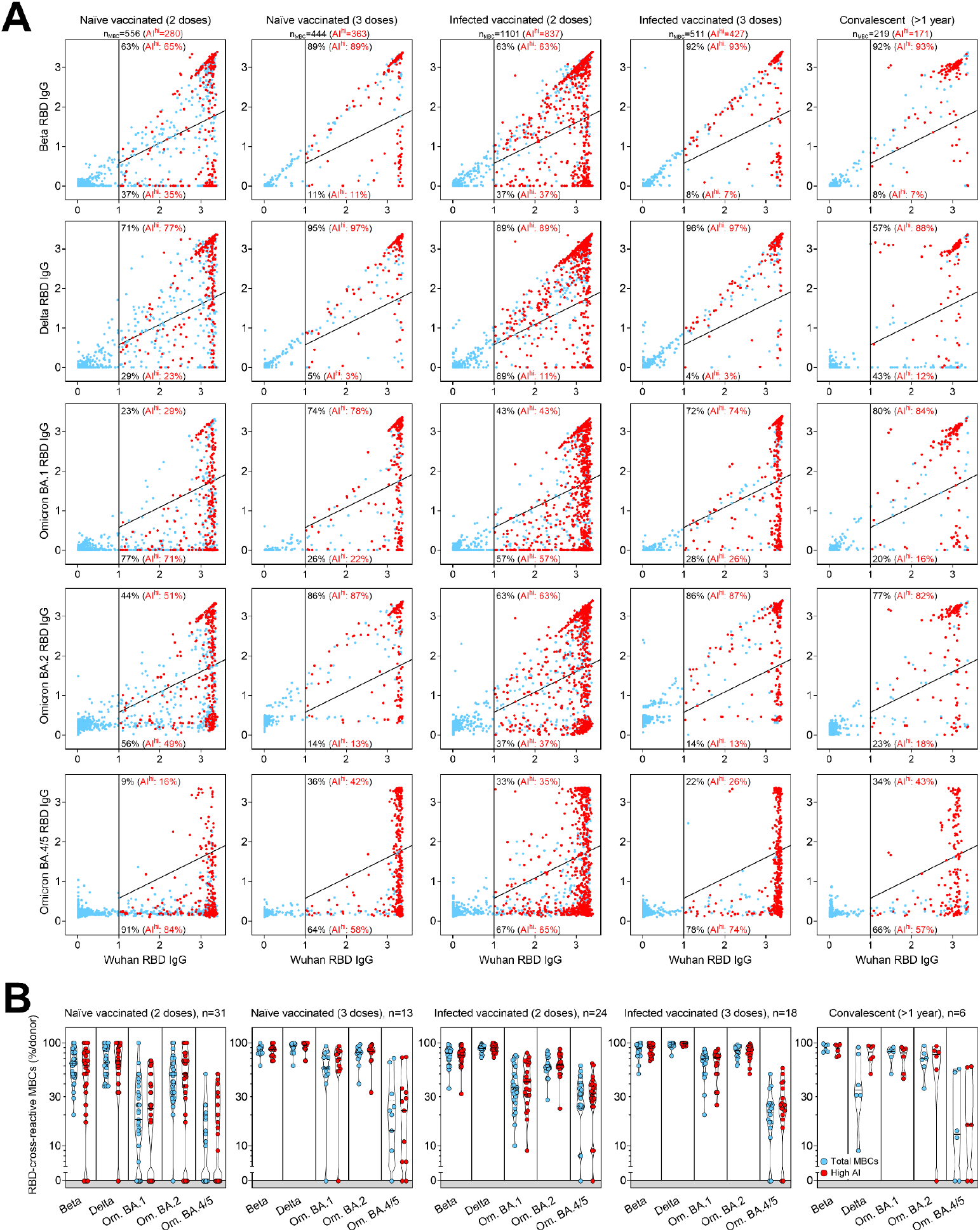
Resilience to viral escape of VOC by high-avidity MBCs primed by SARS-CoV-2 infection and/or vaccination. (A) Cumulative MBC cross-reactivity between RBD from Wuhan SARS-CoV-2 and Beta, Delta, Omicron BA.1, BA.2 and BA.4/5 VOC. Shown are average OD values as measured by ELISA with blank subtracted from n = 2 replicates of 2976, 1248, 2304, 1728 and 576 MBC cultures analyzed from 31 and 13 naïve donors, 24 and 18 infected donors after receiving two and three vaccine doses and from 6 convalescent donors at 376-469 days from symptom onset. RBD-specific MBCs showing high avidity index (AI>80%) are shown in red. Numbers of total and high-avidity RBD-specific MBCs are indicated in the top-left quadrants. Cumulative fractions of total and high-avidity RBD-specific MBCs maintaining or losing binding to the VOC RBD are indicated as percentage in the top-right and bottom-right quadrants. (B) Individual fractions of total and high-avidity SARS-CoV-2 RBD-cross-reactive MBCs that maintain binding with the RBDs of different VOC in convalescent and vaccinated donors. Numbers on top indicate the donors analyzed for the different VOC.

Taken together, these findings indicate that long-lasting affinity maturation upon infection by Wuhan-Hu-1 SARS-CoV-2 and multiple vaccinations can drive the development of MBCs with greater resilience to viral escape.

## DISCUSSION

Kinetics of serum- and MBC-derived antibodies to SARS-CoV-2 after infection and vaccination have been extensively described (Dan *et al*., 2021; Gaebler *et al*., 2021; Goel *et al*., 2021a; Goel *et al*., 2021b; Rodda *et al*., 2021; Roltgen *et al*., 2020; Sokal *et al*., 2021b; Walls *et al*., 2022). In this study, we provide an in depth characterization of the maturation of the memory B cell response to SARS-CoV-2, supporting evidence of how the MBC repertoire is shaped to broaden recall responses to other betacoronaviruses and future variants of concern. Compared to flow-cytometry-based methods (Dan *et al*., 2021; Goel *et al*., 2021a; Rodda *et al*., 2021; Sokal *et al*., 2021b; Wang et al., 2021b), the antigen-specific memory B cell repertoire analysis (AMBRA) has the advantage of analyzing very large numbers of MBCs at the single-cell level, thus allowing unbiased direct comparisons of multiple specificities and functional properties of MBC-derived antibodies.

Documenting the reciprocal kinetics of serum antibodies and MBCs to SARS-CoV-2 antigens illustrates a fundamental aspect of the antibody response. While serum antibodies produced by the first wave of short-lived plasma cells decline over time, MBCs increase in numbers reaching up to 20% of total IgG MBCs in a few months after SARS-CoV-2 infection before their frequencies stabilize. Importantly, this time-dependent increase of MBCs is accompanied by affinity maturation and breadth expansion. As a consequence, while serum antibodies decline, the immune system builds up the capacity to mount a very potent secondary memory response. Accordingly, high numbers of MBCs and breadth against VOC are characteristic of donors who had hybrid immunity due to infection followed by vaccination (Crotty, 2021; Rodda et al., 2022). Conversely, naïve donors require a longer time and multiple immunizations to develop an MBC response of magnitude and breadth, which are comparable to those of infected individuals (Goel et al., 2022; Muecksch et al., 2022). The increased avidity resulting from multiple antigenic stimulations may therefore contribute to broaden the reactivity towards heterologous sarbecoviruses as well as to generate resilience to new VOC (Goel *et al*., 2022; Sokal et al., 2021a; Stamatatos et al., 2021), including Omicron sublineages. This is consistent with the notion that cross-reactive MBCs are primarily induced by repeated antigenic stimulations leading to epitope spread and affinity maturation, which are fundamental for an effective and long-lasting recall response to future SARS-CoV-2 variants. However, the less pronounced resilience observed against the recently emerging BA.4 and BA.5 variants suggests that immune escape, even from high-avidity antibodies, may be a major driver for the evolution of Omicron sublineages.

## SUPPLEMENTAL INFORMATION

## ACKNOWLEDGMENTS

We thank the personnel from the EOC Institute of Laboratory Medicine (EOLAB) for taking blood samples and Ms. Sarita Prosperi for logistics support. F.S. is supported by the Helmut Horten Foundation. O.G. is supported by the Swiss Kidney Foundation. This project has been funded in part with federal funds from the NIAID/NIH under Contract No. DP1AI158186 and HHSN272201700059C to D.V., a Pew Biomedical Scholars Award (D.V.), an Investigators in the Pathogenesis of Infectious Disease Awards from the Burroughs Wellcome Fund (D.V.), Fast Grants (D.V.), and the Natural Sciences and Engineering Research Council of Canada (M.M.). D.V. is an Investigator of the Howard Hughes Medical Institute.

## AUTHOR CONTRIBUTIONS

Experiment design: R.M., J.B., C.S-F., D.C. and L.Pi.

Donors’ recruitment and sample collection: L.Pe., T.T., V.L., M.T., A.R., M.B., A.F.P., C.G., P.F., A.C. and O.G.

Sample processing: R.M., J.B., C.S.-F., F.M., A.C., J.S.L.

Experimental assays: R.M., J.B., C.S.-F., F.M.

Protein expression and purification: K.C., N.S., G.L., C.S., E.C., A.C.W., M.M., M.A.T., J.E.B., E.A.D.J., J.R.D., N.C.

Data analysis: R.M., J.B., C.S.-F., I.B., D.V., A.T., D.C. and L.Pi.

Manuscript writing: C.H.-D., A.A., H.W.V., F.S., D.V., A.L., D.C. and L.Pi.

Supervision: C.G., O.G., A.T., H.W.V., F.S., D.V., A.L., D.C. and L.Pi.

## DECLARATION OF INTERESTS

R.M., J.B., C.S.-F., I.B., F.M., K.C., N.S., G.L., C.S., E.C., E.A.D.J., J.R.D., N.C., A.T., H.W.V., A.L., D.C. and L.Pi. are employees of Vir Biotechnology Inc. and may hold shares in Vir Biotechnology Inc. C.G. is an external scientific consultant to Humabs BioMed SA. Some of the work done in the Veesler laboratory was partially funded by a sponsored research agreement from Vir Biotechnology, Inc. The other authors declare no competing interests.

## STAR METHODS

### KEY RESOURCES TABLE

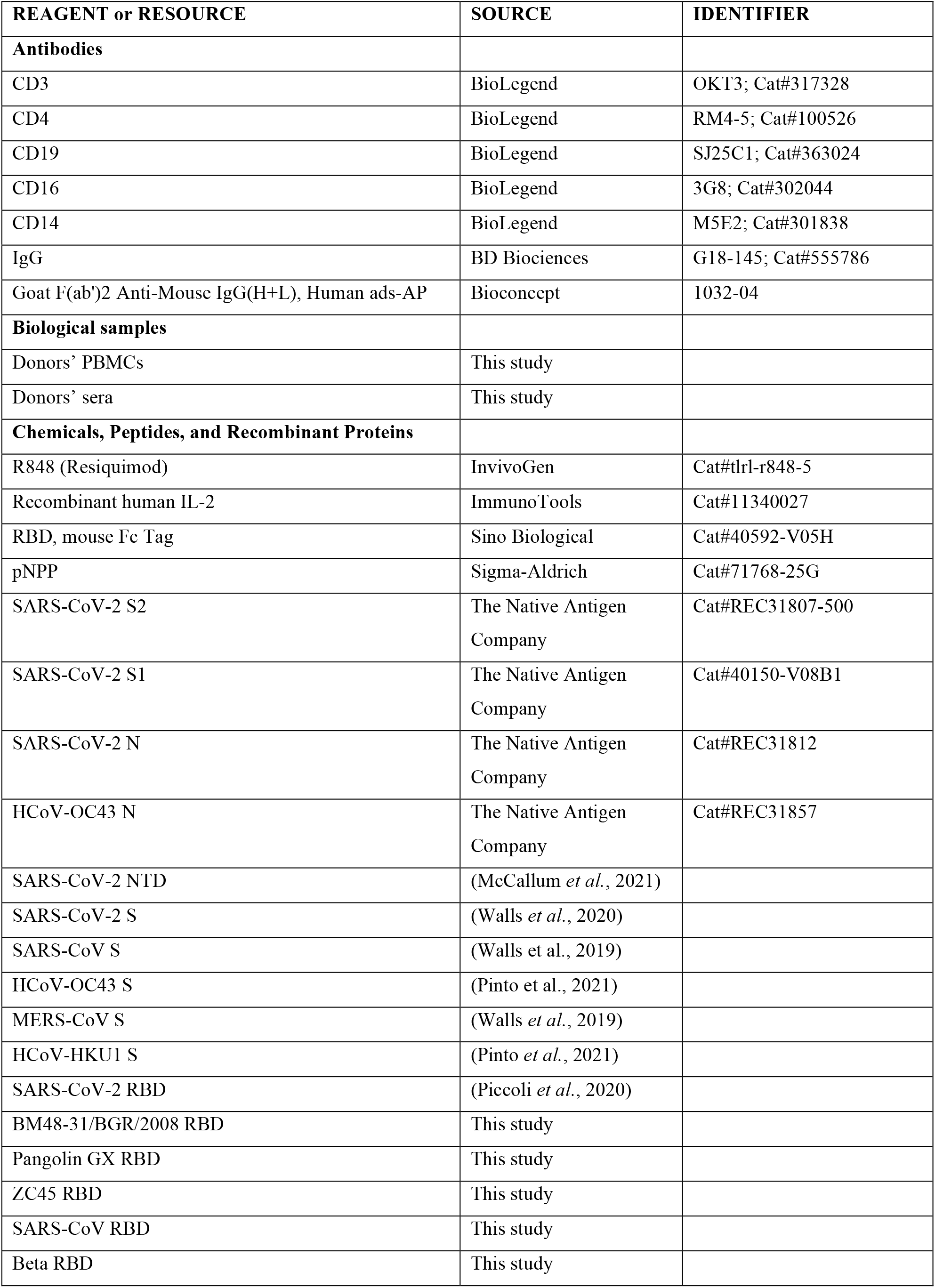
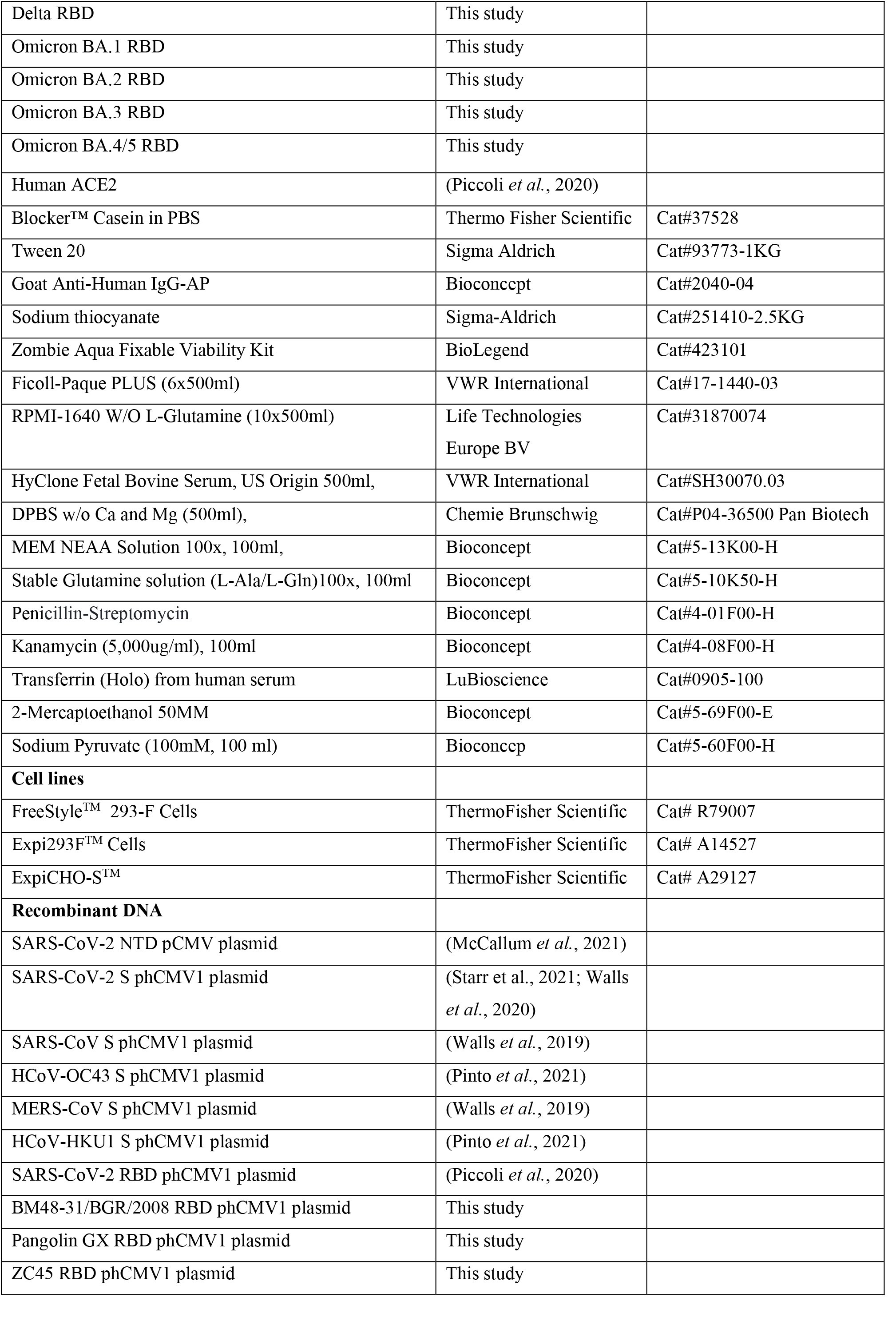
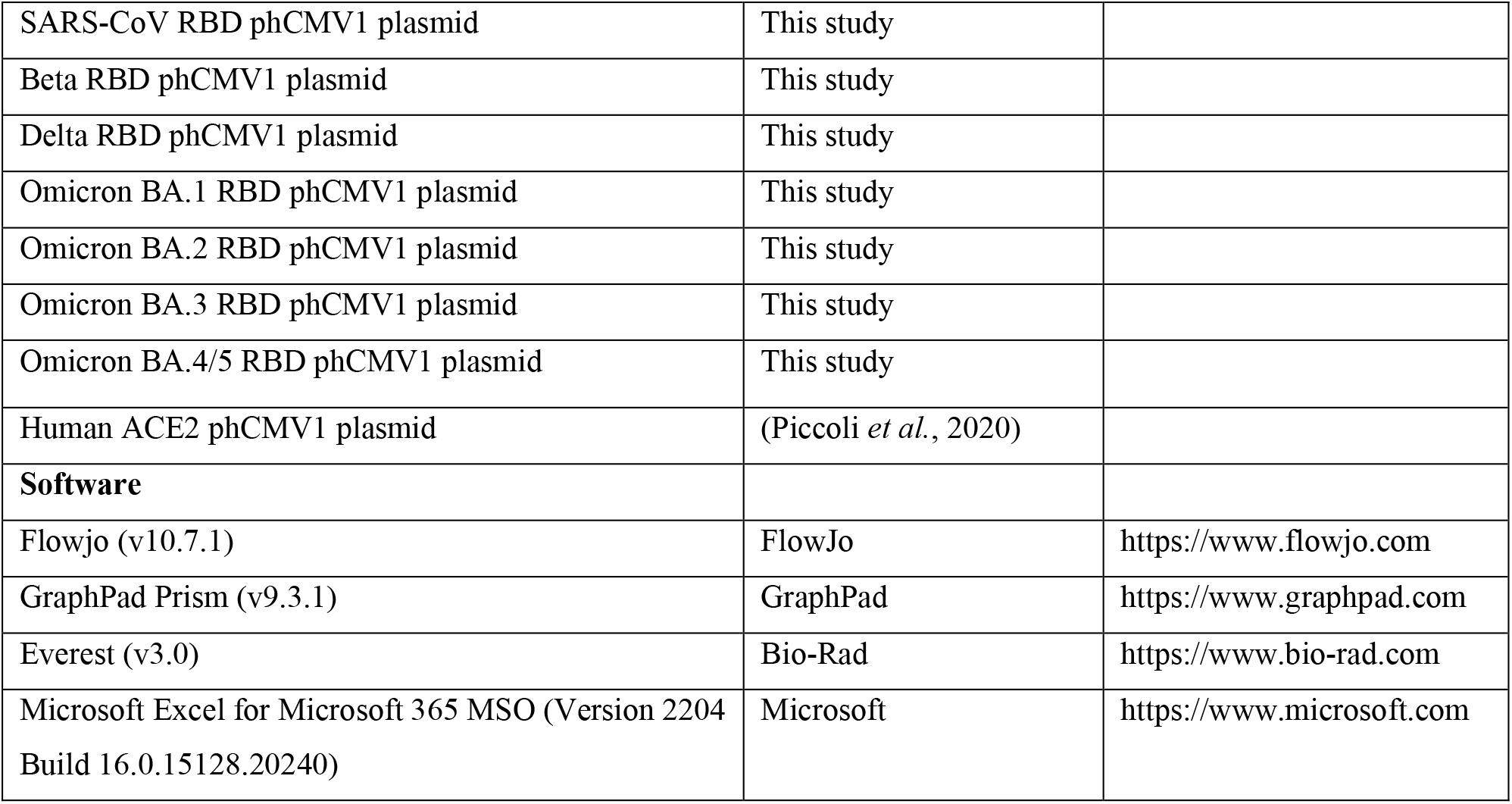

## RESOURCE AVAILABILITY

### Lead contact

Further information and requests for resources and reagents should be directed to and will be fulfilled by the Lead Contact, Luca Piccoli.

### Materials availability

Materials generated in this study will be made available on request and may require a material transfer agreement.

### Data and code availability

Data and code generated in this study will be made available on request and may require a material transfer agreement.

## EXPERIMENTAL MODEL AND SUBJECT DETAILS

### Cell lines

Cell lines used in this study were obtained from ThermoFisher Scientific (FreeStyle™ 293-F Cells, Expi293F™ Cells and ExpiCHO-S™).

### Study participants and ethics statement

Samples were obtained from 3 cohorts of Wuhan SARS-CoV-2-infected individuals, 1 cohort of Alpha SARS-CoV-2-infected individuals and 2 cohorts of individuals vaccinated with Pfizer/BioNtech BNT162b2 mRNA COVID-19 vaccine under study protocols approved by the local Institutional Review Boards (Canton Ticino Ethics Committee, Switzerland and the Ethical committee of Luigi Sacco Hospital, Milan, Italy). COVID-19 was diagnosed by PCR with primers specific for the detection of Wuhan or Alpha SARS-CoV-2 in nasal swaps. All donors provided written informed consent for the use of blood and blood components (such as PBMCs, sera or plasma) and were recruited at hospitals or as outpatients. Based on their availability, participants were enrolled and allocated to either single blood draws or longitudinal follow-up.

## METHOD DETAILS

### Isolation of peripheral blood mononuclear cells (PBMCs), plasma and sera

PBMCs and plasma were isolated from blood draw performed using tubes or syringes pre-filled with heparin or sodium EDTA, followed by Ficoll-Paque PLUS (6×500ml) (VWR International, 17-1440-03) density gradient centrifugation. Sera were obtained from blood collected using tubes containing clot activator, followed by centrifugation. PBMCs, plasma and sera were stored in liquid nitrogen and -80°C freezers until use, respectively.

### Immunophenotyping

PBMCs were thawed and washed twice with RPMI-1640 W/O L-Glutamine (10×500ml) (Life Technologies Europe BV, 31870074) 10% HyClone Fetal Bovine Serum, US Origin 500ml (VWR International, SH30070.03), and incubated in the same medium for 2 h at 37°C. Live PBMCs were counted post thawing and seeded at 1 million into round-bottom 96-well plates (Corning, 3799). PBMCs were stained with LIFE/DEAD marker (Zombie Aqua Fixable Viability Kit, BioLegend 423101) in Dulbecco’s phosphate-buffered saline (DPBS) w/o Ca and Mg (500ml), (Chemie Brunschwig, P04-36500 Pan Biotech) for 30’ at RT, washed in MACS buffer (PBS 2% HyClone, 2 mM EDTA), and stained with antibodies to CD3, CD4, CD19, CD16, CD14, (BioLegend), IgG (BD Bioscience) (Key Resources Table) for 30’ at 4°C. Cells were then washed and resuspended in MACS buffer for data acquisition at ZE5 cytometer (Bio-Rad). Data were analysed with FlowJo software.

### Memory B cell culture

Replicate cultures of total PBMCs were set at different cell densities (10,000-30,000 cells/culture) in 96 U-bottom plates (Corning, 3799). Cells were cultured at 37°C in RPMI 10% Hyclone, 1% Stable Glutamine, 1% Sodium Pyruvate, 1% MEM NEAA, 1% Pen-Strep, 1% Kanamycin, 30 μg/ml Transferrin Holo, 50 μM 2-Mercaptoethanol (50 mM), and stimulated with 2.5 μg/mL R848 (Invivogen, tlrl-r848-5) and 1,000 U/mL human recombinant IL-2 (ImmunoTools, 11340027). Supernatants were harvested after 10 days.

### Plasmid design

The SARS-CoV-2, SARS-CoV, MERS-CoV, HCoV-HKU1 and HCoV-OC43 prefusion S and the SARS-CoV-2 postfusion S ectodomains were synthetized by Genscript or GeneArt and cloned in the phCMV1 vector, as previously described (Bowen *et al*., 2021; Lempp et al., 2021; Pallesen et al., 2017; Pinto *et al*., 2021; Starr *et al*., 2021; Walls *et al*., 2020; Walls *et al*., 2019). The Wuhan-Hu-1 SARS-CoV-2 RBD plasmid, which encodes for the S residues 328-531, was synthetized by Genscript and cloned in the phCMV1 vector, as previously described (Piccoli *et al*., 2020). Plasmids encoding for RBDs of different sarbecoviruses were synthetized by Genscript and cloned in the phCMV1 vector. Plasmids encoding for the RBD of SARS-CoV-2 Beta, Delta, Omicron BA.1, BA.2, BA3 and BA.4/5 variants were generated by overlap PCR (Collier *et al*., 2021). The SARS-CoV-2 NTD plasmid, which encodes for the S residues 14-307, was synthetized by GeneArt and cloned in the pCMV vector, as previously described (McCallum *et al*., 2021).

### Recombinant glycoprotein production

All SARS-CoV-2 S ectodomains were produced in 500-mL cultures of FreeStyle™ 293-F cells (ThermoFisher Scientific) grown in suspension using FreeStyle 293 expression medium (ThermoFisher Scientific) at 37°C in a humidified 8% CO2 incubator rotating at 130 r.p.m. Cells grown to a density of 2.5 million cells per mL were transfected using PEI (9 μg/mL) or 293fectin and respective plasmids, and cultivated for 3-4 days. The supernatant was harvested and, for some productions, cells were resuspended for another three days, yielding two harvests. SARS-CoV-2 S ectodomain was purified from clarified supernatants using a 5-mL C-tag affinity matrix column (Thermo-Fischer). SARS-CoV, MERS-CoV and HCoV-HKU1 S ectodomains were purified using a cobalt affinity column (Cytiva, HiTrap TALON crude). HCoV-OC43 S ectodomain was purified using a 1-ml StrepTrap column (GE Healthcare). All purified proteins were then concentrated using a 100 kDa centrifugal filter (Amicon Ultra 0.5 mL centrifugal filters, MilliporeSigma). Concentrated SARS-CoV-2 S was further purified by a sizing step, using a Superose 6 Increase 10/300 GL column (Cytiva) with 50 mM Tris pH 8, 200 mM NaCl as a running buffer. Peak fractions corresponding to homogeneous spike trimer were pooled. All the proteins were flash frozen in liquid nitrogen and stored for further usage at -80°C.

Postfusion SARS-CoV-2 S, all RBDs and the NTD were produced in Expi293F™ Cells (ThermoFisher Scientific) grown in suspension using Expi293™ Expression Medium (ThermoFisher Scientific) at 37°C in a humidified 8% CO2 incubator rotating at 130 r.p.m. Cells grown to a density of 3 million per mL were transfected using the respective plasmids with the ExpiFectamine™ 293 Transfection Kit (ThermoFisher Scientific) and cultivated for 5 days. SARS-CoV-2 S (used to prepare postfusion S) was purified using a nickel HisTrap HP affinity column (Cytiva) and then incubated with 1:1 w/w S2×58-Fab (Starr *et al*., 2021) and 10 ug/mL trypsin for one hour at 37°C before size exclusion on a Superose 6 Increase column (Cytiva). Supernatants containing RBDs were harvested five days after transfection, equilibrated with 0.1 M Tris-HCl, 0.15 M NaCl, 10 mM EDTA, pH 8.0 and supplemented with a biotin blocking solution (IBA Lifesciences). RBDs were purified by affinity chromatography on a Strep-Trap HP 5 ml column followed by elution with 50 mM biotin and buffer exchange into PBS. The NTD domain was purified from clarified supernatants using 2 ml of cobalt resin (Takara Bio TALON), washing with 50 column volumes of 20 mM HEPES-HCl pH 8.0 and 150 mM NaCl and eluted with 600 mM imidazole. Purified protein was concentrated using a 30 kDa centrifugal filter (Amicon Ultra 0.5 mL centrifugal filters, MilliporeSigma), the imidazole was washed away by consecutive dilutions in the centrifugal filter unit with 20 mM HEPES-HCl pH 8.0 and 150 mM NaCl, and finally concentrated to 20 mg/ml and flash frozen. Recombinant human ACE2 was expressed in Expi293F™ or ExpiCHO-S™ cells transiently transfected with a plasmid encoding for ACE2 residues 19-615, as previously described (Piccoli *et al*., 2020).). Supernatant was collected 6-8 days after transfection, supplemented with buffer to a final concentration of 80 mM Tris-HCl pH 8.0, 100 mM NaCl, and then incubated with BioLock (IBA GmbH) solution. ACE2 was purified using StrepTrap High Performance columns (Cytiva) followed by isolation of the monomeric ACE2 by size exclusion chromatography using a Superdex 200 Increase 10/300 GL column (Cytiva) pre-equilibrated in PBS or 20 mM Tris-HCl pH 7.5, 150 mM NaCl.

### Enzyme-linked immunosorbent assay (ELISA)

Spectraplate-384 with high protein binding treatment (custom made from Perkin Elmer) were coated overnight at 4°C with 1 μg/ml of SARS-CoV-2 S (produced in house), SARS-CoV-2 S2 (The Native Antigen Company, REC31807-500), S1 (The Native Antigen Company, 40150-V08B1), NTD (produced in house), N (The Native Antigen Company, REC31812), SARS-CoV S (produced in house), MERS-CoV S (produced in house), HCoV-HKU1 S (produced in house), HCoV-OC43 S (produced in house), HCoV-OC43 N (The Native Antigen Company, REC31857), RBD SARS-CoV-2 (produced in house), RBD SARS-CoV (produced in house), RBD PangolinGX (produced in house), RBD ZC45 (produced in house), RBD BM48-31/BGR/2008 (produced in house), Beta RBD (produced in house), Delta RBD (produced in house), Omicron BA.1, BA.2 and BA.3 RBD (produced in house) in PBS pH 7.2 or PBS alone as control. Plates were subsequently blocked with Blocker Casein (1%) in PBS (Thermo Fisher Scientific, 37528) supplemented with 0.05% Tween 20 (Sigma Aldrich, 93773-1KG). The coated plates were incubated with diluted B cell supernatant for 1 h at RT. Plates were washed with PBS containing 0.05 % Tween20 (PBS-T), and binding was revealed using secondary goat anti-human IgG-AP (Southern Biotech, 2040-04). After washing, pNPP substrate (Sigma-Aldrich, 71768-25G) was added and plates were read at 405 nm after 1 h or 30’. For chaotropic ELISA, after incubation with B-cell supernatants, plates were washed and incubated with 1 M solution of sodium thiocyanate (NaSCN) (Sigma-Aldrich, 251410-2.5KG) for 1 h. Avidity Index was calculated as the ratio (%) between the ED50 in presence and the ED50 in absence of NaSCN.

### Blockade of RBD binding to human ACE2

Plasma or memory B cell culture supernatants were diluted in PBS and mixed with SARS-CoV-2 RBD mouse Fc-tagged antigen (Sino Biological, 40592-V05H, final concentration 20 ng/ml) and incubated for 30 min at 37°C. The mix was added for 30 min to ELISA 384-well plates (NUNC, P6366-1CS) pre-coated overnight at 4°C with 4 μg/ml human ACE2 (produced in house) in PBS. Plates were washed with PBS containing 0.05 % Tween20 (PBS-T), and RBD binding was revealed using secondary goat anti-mouse IgG-AP (Southern Biotech, 1032-04). After washing, pNPP substrate (Sigma-Aldrich, 71768-25G) was added and plates were read at 405 nm after 1h. The percentage of inhibition was calculated as follow: (1− (OD sample− OD neg ctr)/(OD pos ctr− OD neg ctr)]) × 100.

## QUANTIFICATION AND STATISTICAL ANALYSIS

Data management and statistical analysis were carried out by in-house software based on PostgreSQL and Scala (Odersky et al., 2004). Positive cultures of antigen-specific MBCs were identified from those showing OD values >0.5 by ELISA. This cut-off was determined from 3 times the average OD of pre-pandemic controls. The frequency of B cells precursors specific for a given antigen was calculated assuming a Poisson distribution with the following equation: % of antigen-specific MBCs = -100*(LN(number of negative wells/number of total seeded wells))/number of IgG^+^ MBCs per well. Other statistical and data analyses were performed using GraphPad Prism (v9.3.1) and Microsoft Excel for Microsoft 365 MSO (Version 2204 Build 16.0.15128.20240). Nonparametric Kruskal-Wallis test was used to analyze statistical differences between groups analyzed. Correction for multiple comparison was performed with Dunn’s test. Statistical significance was defined as p <0.05. ED50 values were determined by non-linear regression analysis (log(agonist) versus response - Variable slope (four parameters)). Variation of frequencies and serum titers or avidity over time was determined by one-phase association or decay kinetics models from all the non-null values of each sample.

