## Supplemental Table S1 for "Maturation of SARS-CoV-2 Spike-specific memory B cells drives resilience to viral escape"

**Table S1. Demographics of study participants**

| | | |
|---|---|---|
| **Wuhan SARS-CoV-2 convalescent** | | |
| Participants | | 64 |
| Sex | Female | 20 |
| | Male | 39 |
| | N/A | 5 |
| Age | Average | 55 |
| | Range | 18-95 |
| COVID-19 diagnosis | | March 2020-November 2020 |
| Days after symptom onset | | 13-271 |
| Hospitalized | | 36 |
| | Clinica Luganese Moncucco | 29 |
| | Luigi Sacco Hospital | 7 |
| Symptomatic | | 28 |
| | Clinica Luganese Moncucco | 12 |
| | Swiss volunteers | 16 |
| Subjects with follow-up samples | | 22 |
| | Hospitalized | 9 |
| | Symptomatic | 13 |
| **Alpha SARS-CoV-2 convalescent** | | |
| Participants | | 13 |
| Sex | Female | 11 |
| | Male | 2 |
| Age | Average | 42 |
| | Range | 25-61 |
| COVID-19 diagnosis | | December 2020-January 2021 |
| Symptoms | No | 2 |
| | Yes | 11 |
| Days after symptom onset | | 30-53 |
| **Vaccinated donors** | | |
| Participants | | 78 |
| Sex | Female | 49 |
| | Male | 29 |
| Age | Average | 42 |
| | Range | 24-67 |
| SARS-CoV-2 | Naïve | 46 |
| (days before vaccination) | Previously infected (53-389, 157 average days) | 32 |
| Vaccine | Pfizer/BioNtech BNT162b2 | |
| Vaccine doses | 1 (12-33, average 16 days) | 78 |
| (sampling time after | 2 (10-21, average 15 days / 125-293, 166 days) | 76 |
| vaccination) | 3 (10-35, average 21 days) | 44 |
| Hospital | Clinica Luganese Moncucco | 29 |
| | Ente Ospedaliero Cantonale | 49 |
| Subjects with a pre-vaccine sample | | 61 |
| Subjects with post-vaccine follow-up samples | | 63 |
