## Supplemental Figures S1-6 for "Maturation of SARS-CoV-2 Spike-specific memory B cells drives resilience to viral escape"

### Figure S1

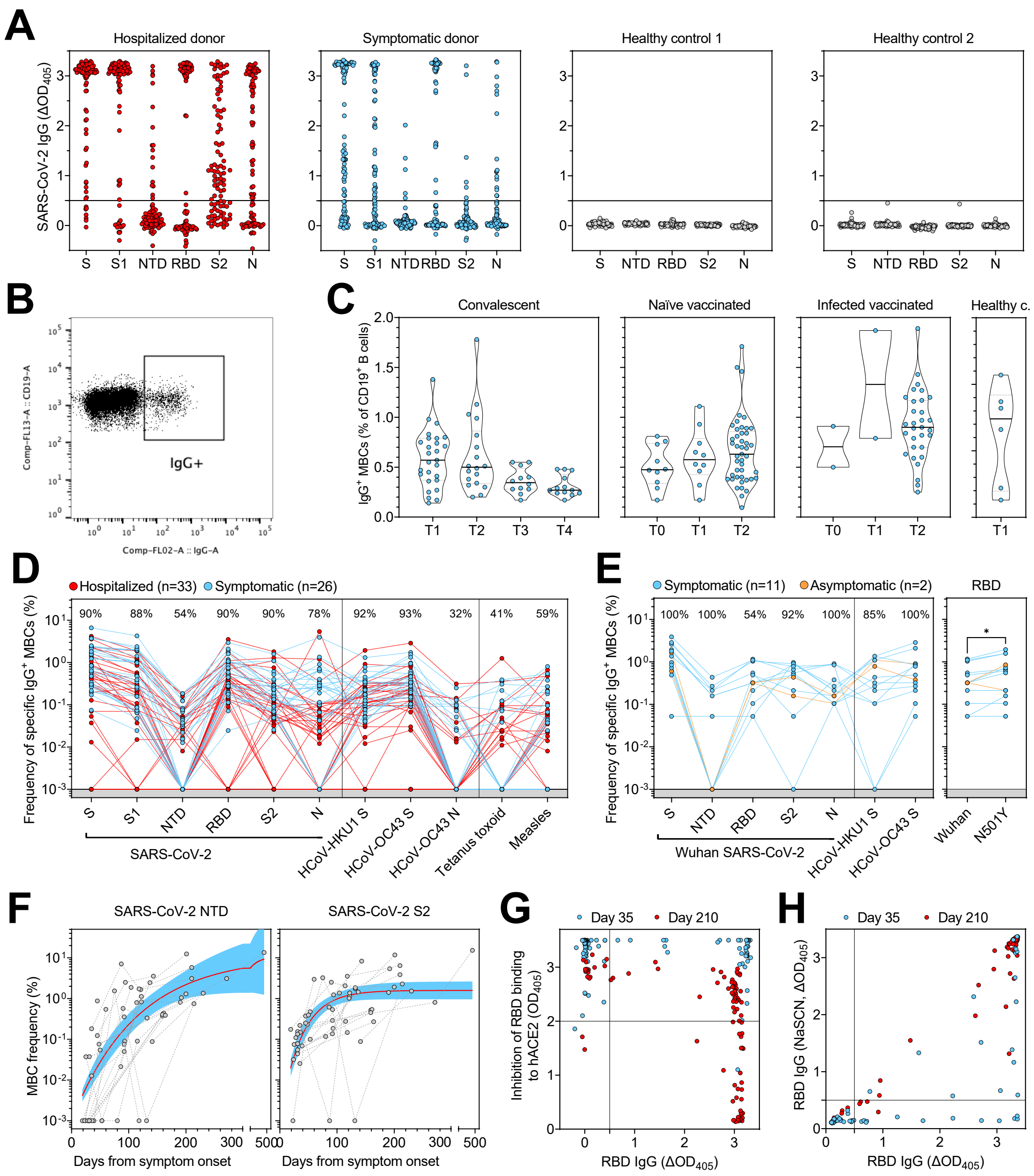

A
Hospitalized donor
Symptomatic donor
Healthy control 1
Healthy control 2
SARS-CoV-2 IgG (ΔOD405)
S S1 NTD RBD S2 N
B
Comp-FL13-A :: CD19-A
Comp-FL02-A :: IgG-A
IgG+
C
Convalescent
Naïve vaccinated
Infected vaccinated
Healthy c.
IgG+ MBCs (% of CD19+ B cells)
T0 T1 T2 T3 T4
D
Hospitalized (n=33)
Symptomatic (n=26)
Frequency of specific IgG+ MBCs (%)
90% 88% 54% 90% 90% 78% 92% 93% 32% 41% 59%
SARS-CoV-2
HCoV-HKU1 S
HCoV-OC43 S
HCoV-OC43 N
Tetanus toxoid
Measles
E
Symptomatic (n=11)
Asymptomatic (n=2)
RBD
100% 100% 54% 92% 100% 85% 100%
Wuhan SARS-CoV-2
Wuhan
N501Y
F
SARS-CoV-2 NTD
SARS-CoV-2 S2
MBC frequency (%)
Days from symptom onset
G
Day 35
Day 210
Inhibition of RBD binding to hACE2 (OD405)
RBD IgG (ΔOD405)
H
RBD IgG (NaSCN, ΔOD405)

**Figure S1. Analysis of frequencies of memory B cells, Related to Figure 1**

(A) Memory B cell reactivity to different Wuhan SARS-CoV-2 antigens from 2 representative donors, one hospitalized (red) and one symptomatic (blue), and 2 healthy controls. Shown are average OD values as measured by ELISA with blank subtracted from n = 2 replicates of 96 MBC cultures analyzed for each donor. S, Spike. Cut-off is shown by a black line at 0.5 OD. RBD, receptor binding domain. NTD, N-terminal domain. N, Nucleoprotein.

(B) Representative scatter density plot of flow cytometry data used to define the frequency of $IgG^+$ MBCs expressed as percentage of total $CD19^+$ MBCs isolated from n = 77 convalescent and 79 vaccinated donors.

(C) Frequency of $IgG^+$ MBCs from n = 76 samples collected at different time points after symptom onset (T1 to T4), before (T0) or after vaccination (T1, after dose 1, and T2, after dose 2).

(D) Frequency of SARS-CoV-2-specific MBCs isolated from 33 hospitalized (red) and 26 symptomatic donors (blue). Shown is the reactivity to antigens of SARS-CoV-2 and other betacoronaviruses (HCoV-HKU1 and HCoV-OC43): Spike (S), S1 domain, N-terminal domain (NTD), receptor-binding domain (RBD), S2 domain, Nucleoprotein (N). Reactivities to Tetanus toxoid and to Measles virus (lysate) are included as controls. Lines connect data from the same donors. Percentages of donors with detectable specific MBCs are indicated above each set of data.

(E) Frequency of SARS-CoV-2-specific MBCs isolated 11 symptomatic (blue) and 2 asymptomatic donors (orange) infected with Alpha SARS-CoV-2. The left panels shows the reactivity to antigens of Wuhan SARS-CoV-2 and other betacoronaviruses (HCoV-HKU1 and HCoV-OC43): S, NTD, RBD, S2 and N. The right panel shows a comparison of frequencies of MBCs specific for Wuhan and N501Y RBD in 13 Alpha SARS-CoV-2-infected individuals. Lines connect data from the same donors. Percentages of donors with detectable specific MBCs are indicated above each set of data. A significant difference is indicated as * ($p < 0.033$).

(F) Frequency of MBCs specific for SARS-CoV-2 NTD and S2 from n = 21 donors followed-up up to 469 days after symptom onset. Frequencies were obtained from the analysis of 5,760 MBC cultures (60 samples, minimum 2 samples per donor). Black dotted lines connect samples from the same donor. A one-phase association kinetics model (red line) was calculated from all the non-null values of each sample. The area within 95% confidence bands is shown in blue.

(G) SARS-CoV-2 RBD binding of IgG MBCs compared to their ability of inhibiting RBD binding to ACE2. Shown are 2 time points from one representative donor. Specific RBD-binding is set for $\Delta OD > 0.5$ and inhibition of ACE2 is set for $OD < 2$.

(H) SARS-CoV-2 RBD binding of IgG MBCs in presence or absence of chaotropic agent NaSCN. Shown are 2 time points from one representative donor.

### Figure S2

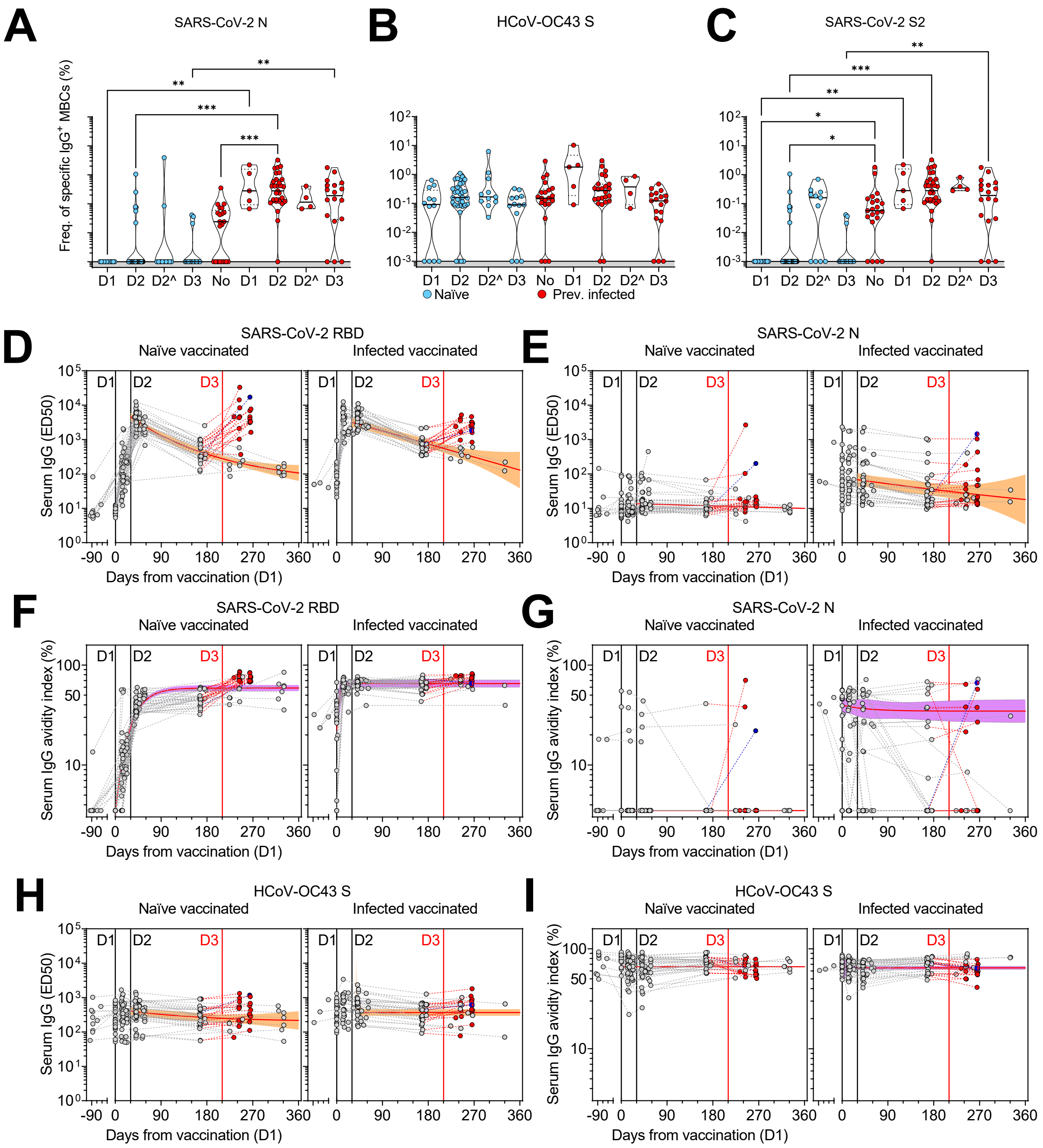
A
SARS-CoV-2 N
Freq. of specific IgG+ MBCs (%)
D1 D2 D2^ D3 No D1 D2 D2^ D3
B
HCoV-OC43 S
Naïve
Prev. infected
C
SARS-CoV-2 S2
D
SARS-CoV-2 RBD
Naïve vaccinated
Infected vaccinated
Serum IgG (ED50)
Days from vaccination (D1)
E
SARS-CoV-2 N
F
SARS-CoV-2 RBD
Serum IgG avidity index (%)
G
SARS-CoV-2 N
H
HCoV-OC43 S
I
HCoV-OC43 S

**Figure S2. Analysis of frequencies of vaccine-induced MBC- and serum-derived antibodies, Related to Figure 2**

(A-C) Frequency of MBCs specific for SARS-COV-2 N (A), HCoV-OC43 S (B) and S2 (C) of n = 12, 45 and 13 naïve donors and 5, 31, and 18 previously infected donors 10-35 days after the first (D1), the second (D2) and the third dose (D3) of Pfizer/BioNtech BNT162b2 mRNA vaccine, respectively. Shown are also 11 naïve and 4 immune donors whose MBCs were isolated 125-293 days after the second dose (D2^). Median frequencies are compared withing donor groups and between respective vaccine doses as well as to a group of n = 21 convalescent donors at 18-30 days after symptom onset. Significant differences are indicated as *** (p-value $< 0.001$); ** ($p < 0.002$), * ($p < 0.033$), ns (non-significant, $p > 0.12$).

(D-E) Serum IgG ED50 titers to SARS-CoV-2 RBD (D) and N (E) of samples collected from n = 47 naïve (left) and 32 immune donors (right) 10-35 days after the first (D1), the second (D2) and the third dose (D3) of Pfizer/BioNtech BNT162b2 mRNA vaccine, respectively. A one-phase decay kinetics model (red line) was calculated from all the non-null values of each sample and the area within 95% confidence bands is shown in orange. 37 samples collected from individuals who received a third dose (red) or had a second SARS-CoV-2 infection (blue) were excluded from the decay analysis.

(F-G) Serum IgG avidity indexes to SARS-CoV-2 RBD (F) and N (G) of the same samples shown in panels D-E. A one-phase association kinetics model (red line) was calculated from all the non-null values of each sample and the area within 95% confidence bands is shown in violet.

(H-1) Serum IgG ED50 titers (H) and avidity indexes (I) to HCoV-OC43.

### Figure S3

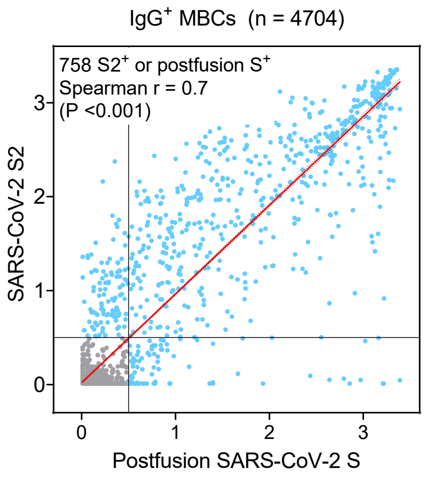


**Figure S3. SARS-CoV-2 S2 is a proxy for the postfusion conformation of the Spike, Related to Figure 3**

Correlation analysis of the cross-reactivity of MBC specific for SARS-CoV-2 S2 and/or the S stabilized in its postfusion conformation. Shown are average OD values as measured by ELISA with blank subtracted from n = 2 replicates of 4704 MBC cultures of which 758 are S2 and/or postfusion S-specific.

### Figure S4

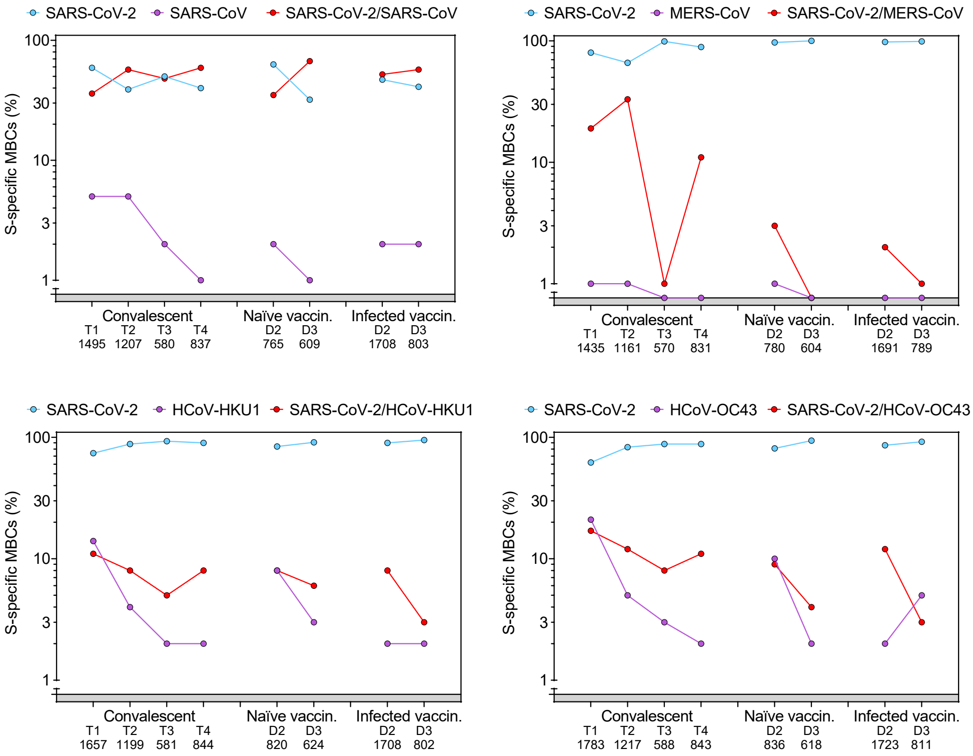


**Figure S4. Follow-up analysis of cross-reactivity to betacoronaviruses of MBCs, Related to Figure 4**

Cumulative MBC single reactivity and cross-reactivity between the S of SARS-CoV-2 and the four betacoronaviruses SARS-CoV, MERS-CoV, HCoV-HKU1 and HCoV-OC43 in convalescent donors analyzed at multiple time points after symptom onset (T1, T2, T3 and T4) and in naïve and infected donors after two (D2) and three (D3) vaccine doses. Number of S-specific MBCs analyzed are indicated below the x-axis.

### Figure S5

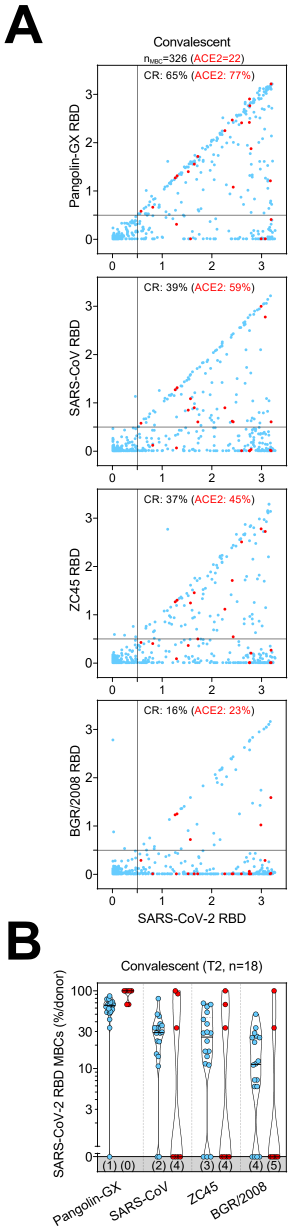


**Figure S5. Cross-reactivity to sarbecoviruses of MBCs primed by SARS-CoV-2 infection at later time points, Related to Figure 5**

(A) Cumulative MBC cross-reactivity between SARS-CoV-2 RBD and four sarbecoviruses representative of clades 1a (SARS-CoV), 1b (Pangolin Guangxi), 2 (ZC45) and 3 (BM48-31/BGR/2008). Shown are average OD values as measured by ELISA with blank subtracted from n = 2 replicates of 1728 MBC cultures analyzed from 18 convalescent donors' samples collected up to 141 days after symptom onset. RBD-specific MBCs showing inhibition of binding to ACE2 are shown in red. Cumulative fractions of total and ACE2-inhibiting RBD- cross-reactive (CR) MBCs are indicated as percentage in the top-right quadrant.

(B) Individual fractions of SARS-CoV-2 RBD-specific MBCs that cross-react with the four representative sarbecoviruses in convalescent and vaccinated donors. Numbers in brackets indicate the donors with MBCs showing no cross-reactivity for the respective sarbecovirus RBD.

### Figure S6

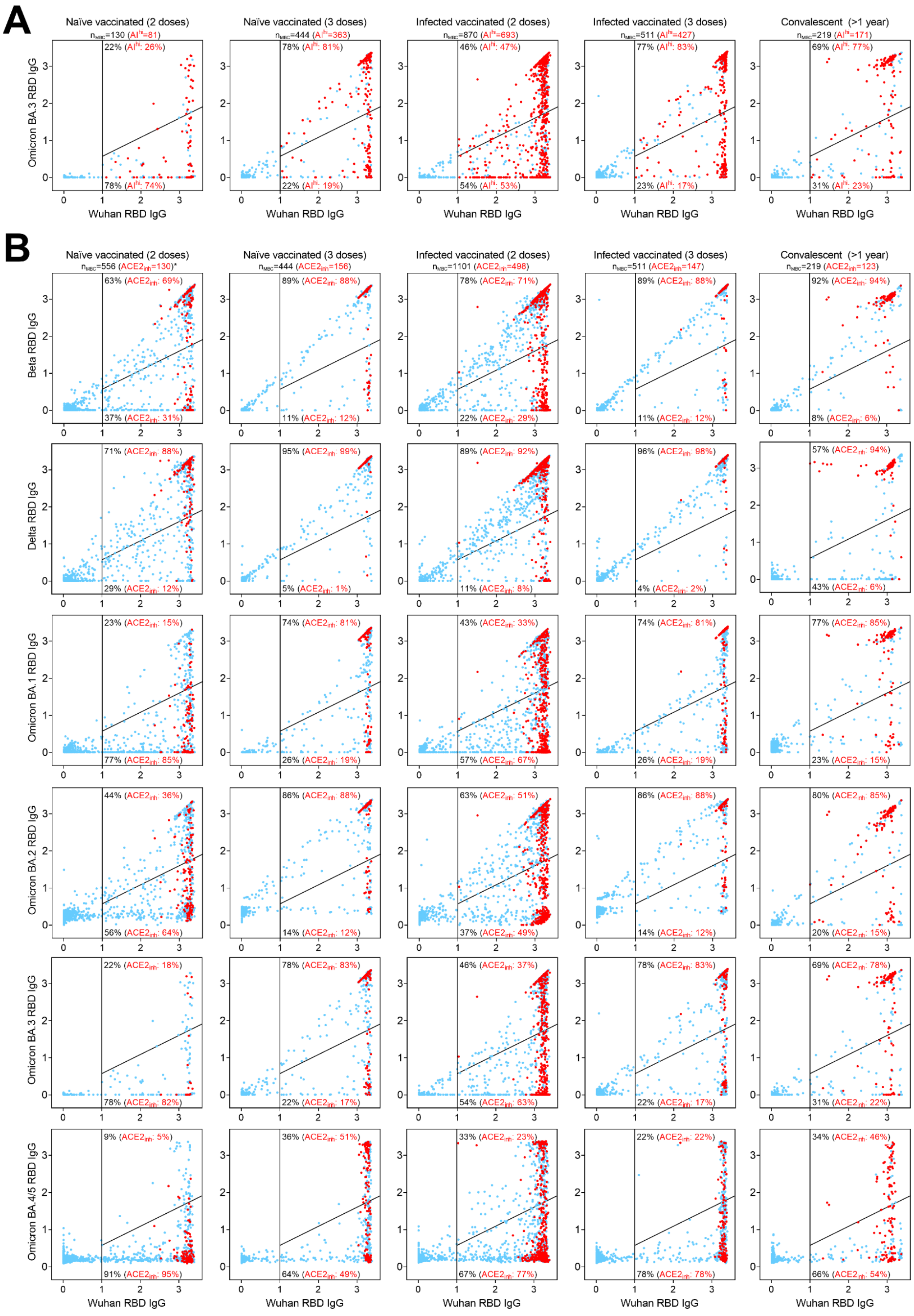


**Figure S6. Analysis of resilience of ACE2-inhibiting MBCs primed by SARS-CoV-2 infection and/or vaccination, Related to Figure 6**

(A) Cumulative MBC cross-reactivity between RBD from Wuhan SARS-CoV-2 and Omicron BA.3 VOC.

(B) Cumulative MBC cross-reactivity between RBD from Wuhan SARS-CoV-2 and Beta, Delta, Omicron BA.1, BA.2, BA.3 and BA.4/5 VOC. RBD-specific MBCs showing inhibition of binding to ACE2 are shown in red. *Cross-reactivity analyzed for BA.3 in naïve vaccinated (2 doses) donors: $n_{MBC}$=130 ($ACE2_{inh}$=22). Cumulative fractions of total and ACE2-inhibiting RBD-specific MBCs maintaining or losing binding to the VOC RBD are indicated as percentage in the top-right and bottom-right quadrants.
